# The role of mutation bias in adaptive molecular evolution: insights from convergent changes in protein function

**DOI:** 10.1101/580175

**Authors:** Jay F. Storz, Chandrasekhar Natarajan, Anthony V. Signore, Christopher C. Witt, David M. McCandlish, Arlin Stoltzfus

## Abstract

An underexplored question in evolutionary genetics concerns the extent to which mutational bias in the production of genetic variation influences outcomes and pathways of adaptive molecular evolution. In the genomes of at least some vertebrate taxa, an important form of mutation bias involves changes at CpG dinucleotides: If the DNA nucleotide cytosine (C) is immediately 5’ to guanine (G) on the same coding strand, then – depending on methylation status – point mutations at both sites occur at an elevated rate relative to mutations at non-CpG sites. Here we examine experimental data from case studies in which it has been possible to identify the causative substitutions that are responsible for adaptive changes in the functional properties of vertebrate hemoglobin (Hb). Specifically, we examine the molecular basis of convergent increases in Hb-O_2_ affinity in high-altitude birds. Using a data set of experimentally verified, affinity-enhancing mutations in the Hbs of highland avian taxa, we tested whether causative changes are enriched for mutations at CpG dinucleotides relative to the frequency of CpG mutations among all possible missense mutations. The tests revealed that a disproportionate number of causative amino acid replacements were attributable to CpG mutations, suggesting that mutation bias can influence outcomes of molecular adaptation.

## 1. Introduction

A question of enduring interest in EvoDevo research concerns the extent to which the biased production of variation during development influences directional trends and patterns of convergence in morphological evolution [1-4]. At the molecular sequence level, an analogous question concerns the extent to which mutational bias in the production of genetic variation influences pathways of molecular evolution [5, 6]. Mutation bias is known to exert an important influence on patterns of neutral molecular evolution [7], but an emerging understanding of the implications of mutation as an introduction process suggest that it may be an important orienting factor in adaptive evolution as well [8, 9]. The Modern Synthesis originally included a strong position on the importance of recombination, and did not attach much importance to the role of mutation bias [10-13]. When the process of evolution is defined in terms of shifts in the frequencies of pre-existing alleles, recombination serves as the main source of new genetic variation and mutation rates must be on the order of selection coefficients to exert an appreciable influence on the direction of adaptive change [14].

An alternative view is that outcomes and pathways of evolution may be influenced by the rate of mutation even in the presence of selection [5]. For instance, in the simple case of origin-fixation models [8], the substitution rate is given as *K* = 2*N*μλ, where *N* is the size of a diploid population, μ is the per-copy rate of mutation, and λ is the probability of fixation [15]. This model specifies the substitution rate as the product of the rate at which new alleles originate via mutation (2*N*μ) and the probability that they become fixed once they arise (λ). In this type of model, mutation bias in the introduction of variation can produce a bias in substitution rates even when the substitutions are beneficial [8]. Results of several experimental evolution studies suggest that mutation bias can influence trajectories of adaptive protein evolution [9, 16-20], and Stoltzfus and McCandlish [9] also provide evidence for transition-transversion bias among adaptive substitutions that contributed to natural protein evolution.

One especially powerful means of addressing questions about the role of mutation bias in molecular adaptation is to examine convergent changes in protein function that can be traced to specific amino acid substitutions. The experimental identification of causative substitutions provides a means of testing whether such changes are random with respect to different classes of site or different classes of mutational change. In vertebrate genomes, transition:transversion bias results in especially high rates of change from one pyrimidine to another (C↔T) or from one purine to another (G↔A). In the genomes of mammals and birds, the dinucleotide CG – often designated “CpG” – is a hotspot of nucleotide point mutations due to the effect of methylation on damage and repair. In mammals, the mutation rate at CpG sites is elevated 10-fold for transition mutations and several-fold for transversions [21, 22]. A recent mutation-accumulation study in birds confirmed a similar increase in mutation rate at CpG sites [23]. Mammalian and avian genomes both exhibit roughly five-fold depletions of CpG dinucleotides, consistent with elevated rates of mutation at such sites [24].

In studies of hemoglobin (Hb) evolution in high-altitude birds, site-directed mutagenesis experiments have documented three cases in which missense mutations at CpG dinucleotides contributed to derived increases in Hb-O_2_ affinity: 55Ile→Val (I55V) in the β^*A*^-globin gene of Andean house wrens (*Troglodytes aedon*)[25] and parallel 34Ala→Thr substitutions (A34T) in the α^*A*^-globin genes of two different passerine species that are native to the Tibetan Plateau, the ground tit (*Parus humilis* [= *Pseudopodoces humilis*]) and the grey crested tit (*Lophophanes dichrous*)[26]. The patterns documented in these three high-altitude taxa are consistent with a broader pattern of convergence in Hb function in highland birds [26-30], and the direction of character-state change is consistent with theoretical and experimental results which suggest that it is generally beneficial to have an increased Hb-O_2_ affinity under conditions of severe environmental hypoxia [30-32]. Moreover, in the case of the high-altitude house wrens, an analysis of genome-wide patterns of nucleotide polymorphism revealed evidence for altitude-related selection on the missense CpG variant [25], providing additional support for the inferred adaptive significance of the affinity-altering amino acid change.

The documented cases in which CpG mutations have contributed to altitude-related changes in Hb function suggest that mutation bias may influence which mutations are most likely to contribute to molecular adaptation. However, a far more comprehensive and systematic analysis is required to draw firm conclusions. Here we examine existing data from a larger set of case-studies in which it has been possible to identify the causative substitutions that are responsible for convergent increases in Hb-O_2_ affinity in high-altitude birds. Using a data set of experimentally verified, affinity-enhancing mutations in the Hbs of highland avian taxa, we test whether a disproportionate number involve mutations at CpG sites. The results suggest that mutation bias has influenced outcomes of molecular adaptation.

## 2. Methods

### (a) Sampling design

To test for convergent changes in Hb-O_2_ affinity, we conducted phylogenetically independent comparisons involving 70 avian taxa representing 35 matched pairs of high-versus low-altitude species or subspecies (**Fig. 1**). These taxa include ground doves, nightjars, hummingbirds, passerines, and waterfowl [25-29, 33-36]. All pairwise comparisons involved dramatic elevational contrasts; high-altitude taxa native to very high elevations in the Andes, Ethiopian highlands, or Tibetan Plateau (with upper range limits of 3,500 to 5,000 m above sea level) were paired with close relatives that typically occur at or near sea level. Thus, the highland members of each taxon pair have upper elevational range limits that likely exceed the threshold at which an increased Hb-O_2_ affinity becomes physiologically beneficial.

**Fig. 1.**
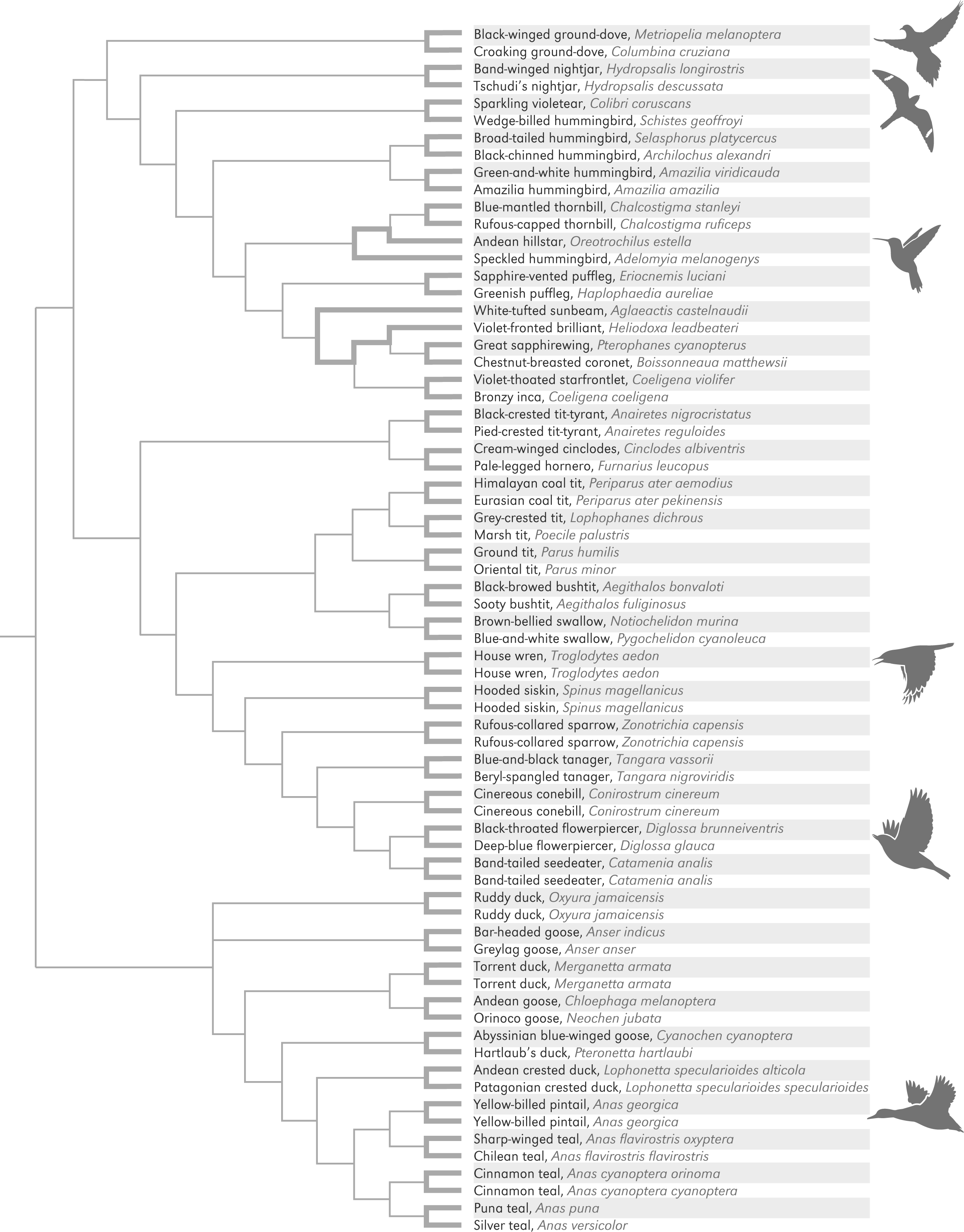
Phylogenetic relationships of 70 avian taxa used in comparative analyses of Hb function. Rows corresponding to high-altitude taxa are shaded. Branches in bold connect pairs of high- and low-altitude taxa that were used to test for a relationship between Hb-O_2_ affinity and native elevation. The 35 pairwise comparisons are phylogenetically independent. For information regarding elevational ranges, see refs. [25-29, 33].

**Fig. 2.**
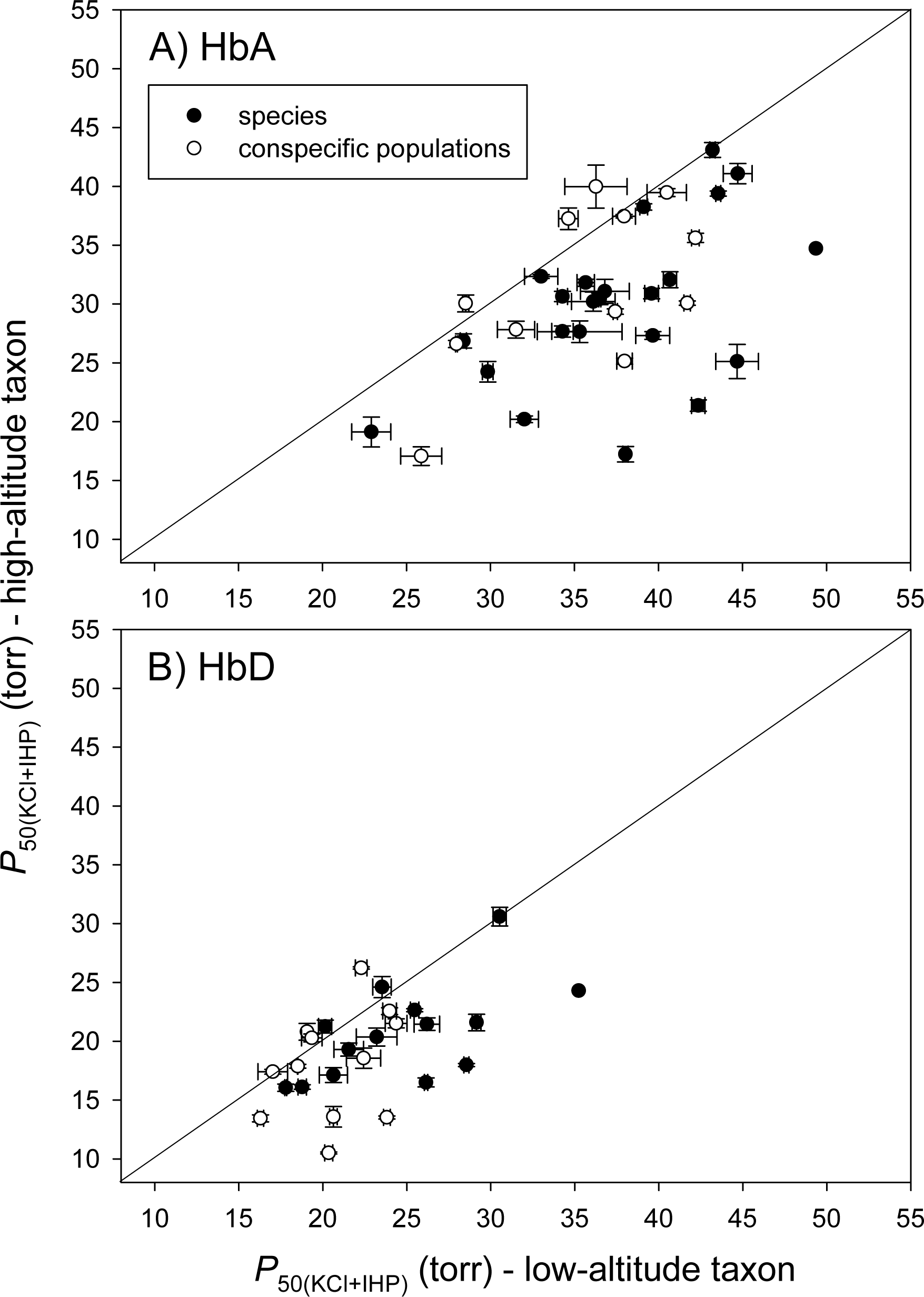
Convergent increases in Hb–O_2_ affinity in high-altitude Andean birds. (A) Plot of *P*_50(KCl+IHP)_ (± SE) for HbA in 35 matched pairs of high- and low-altitude taxa. The pairwise comparisons of Hb–O_2_ affinity involve HbA homologs of closely related species or representative allelic variants from high- and low-altitude populations of the same species. Data points that fall below the diagonal (x = y) denote cases in which the high-altitude member of a given taxon pair possesses a higher Hb–O_2_ affinity (lower *P*_50_). Comparisons involve phylogenetically replicated pairs of taxa, so all data points are statistically independent. (B) Plot of *P*_50(KCl+IHP)_ (± SE) for the minor HbD isoform in a subset of the same taxon pairs in which both members of the pair express HbD. Sample sizes are larger for HbA than for HbD because the two ground dove species (*Metriopelia melanoptera* and *Columbina cruziana*) expressed no trace of HbD, and several hummingbird species expressed HbD at exceedingly low levels [27, 29]. In such cases, sufficient quantities of HbD could not be purified for measures of O_2_ equilibria. Black symbols denote comparisons between species, whereas open symbols denote comparisons between high-versus low-altitude populations of the same species. Data from refs. [26, 28, 29].

### (b) Experimental measurement of Hb function

For each taxon, we used our previously measured O_2_ affinities of purified Hbs in the presence of allosteric cofactors: Cl^-^ ions (added as KCl) and inositol hexaphosphate (IHP). IHP is a chemical analog of inositol pentaphosphate (IPP), which is endogenously produced as a metabolite of oxidative phosphorylation in avian red cells. Using a diffusion-chamber protocol, we measured O_2_ equilibria of purified Hb solutions at pH 7.40, 37 °C in the presence and absence of allosteric cofactors, namely 0.10 M Cl^-^; 0.1 M HEPES (4-(2-hydroxyethyl)-1-piperazineethanesulfonic acid); 0.3 mM heme; and IHP:tetrameric Hb ratio of 2.0 [26, 37-39]. Estimated values of *P*_50_ (the partial pressure of O_2_ at which Hb is 50 % saturated) provide an inverse measure of Hb-O_2_ affinity.

Most bird species express two main isoforms of the Hb heterotetramer (α_2_ β_2_) in adult red blood cells: HbA (the major isoform, with α-chain subunits encoded by the α^*A*^-globin gene) and HbD (the minor isoform, with α-chain subunits encoded by the α^*D*^-globin gene); both isoforms incorporate the same β-chain subunits [39, 40]. In species that expressed both Hb isoforms, we separately measured the O_2_ -binding properties of purified HbA and HbD.

All the experimental data used in the present analysis were published previously [25-29, 33-36], and these papers can be consulted for additional details regarding experimental protocols. For additional methodological details regarding the expression and purification of recombinant Hbs, see Refs. [29, 41-43]. We restrict the analysis to data based on standardized measurements of purified Hbs, so the observed variation in *P*_50_ values is purely genetic, reflecting evolved changes in the amino acid sequences of the α- and β-type subunits of the α_2_β_2_ Hb tetramer. This focus on purified Hbs avoids problems associated with the confounding effects of environmentally induced variation.

### (c) Testing for convergence in Hb function

In comparative analyses of phenotypic variation it is important to account for the fact that trait values from different species are not statistically independent because the sampled species did not evolve independently of one another [44]. Accordingly, we used a paired-lineage design [45] to test for a nonrandom association between Hb-O_2_ affinity and native elevation in the set of 70 avian taxa. The paired-lineage design restricts comparisons to phylogenetically replicated pairs of high- and low-altitude taxa that were chosen so that there is no overlap in evolutionary paths of descent. Since our comparative analysis includes a phylogenetically diverse range of avian taxa, an advantage of this design is that comparisons can be restricted to closely related species by excluding pairs with long paths between them [30].

### (d) Experimental identification of causative mutations

For each pair of high- and low-altitude taxa in which we documented an evolved difference in Hb-O_2_ affinity, we used a combinatorial protein-engineering approach to identify the causative substitution(s). Specifically, we synthesized recombinant Hbs representing the wildtype Hbs of the high- and low-altitude species, and we used ancestral sequence reconstruction to synthesize the recombinant Hb representing the inferred genotype of the species’ most recent common ancestor. By design, our pairs of high- and low-altitude taxa were always very closely related (in many cases representing sister species or nominal subspecies) so the ancestral state estimates were always unambiguous [26, 28, 29, 36]. In cases where the triangulated comparison involving the reconstructed ancestral protein and those from the pair of modern-day descendants confirmed that the highland taxon evolved a derived increase in Hb-O_2_ affinity, we performed site-directed mutagenesis experiments to measure the additive and nonadditve effects of all or nearly all amino acid substitutions that were specific to the highland lineage [25-29, 33, 36]. In the majority of cases, we synthesized combinatorially complete sets of mutant genotypes in order to test each mutation in all possible multi-site combinations. There were two pairs of bird species for which the site-directed mutagenesis experiments were not combinatorially complete: the comparison between *Parus humilis* and *Parus minor* and that between *Lophophanes dichrous* and *Poecile palustris*. In both of these pairwise comparisons, we experimentally tested mutations at a single candidate site so we cannot rule out the possibility that additional, untested substitutions also contributed to the observed differences in Hb-O_2_ affinity between the major HbA isoforms of the high- and low-altitude species.

### (e) Null model for frequency of CpG changes among causative mutations

To assess a role for CpG effects in molecular adaptation, we calculated the expected frequency of adaptive CpG changes for a null model in which CpG status is irrelevant to adaptation, and we compared the observed frequency of adaptive CpG changes to this null expectation. In all cases examined here, outgroup comparisons revealed that the derived, affinity-enhancing amino acid changes were specific to the high-altitude taxa. Using reconstructed ancestral sequences for each pair of taxa, we calculated the null expectation for the frequency of CpG changes for three slightly different models (custom R script, Dryad Digital Repository: https://datadryad.org/review?doi=doi:10.5061/dryad.2256f38).

In particular, following Stoltzfus and McCandlish [9], we considered null models in which all mutationally accessible variants are maintained in the population at sufficiently high frequency that selection deterministically chooses the fittest variant independent of its mutation rate or CpG status. Because we do not know *a priori* which variant will be the fittest, we calculate the expected frequency of CpG-associated changes by assuming that each mutational neighbor has an equal probability of being the most fit. Although the most obvious justification for this kind of calculation is in terms of the population-genetic scenario just described, the same null expectation applies to any scenario in which CpG status (for whatever reason) fails to have any impact on which mutant is chosen from the set of mutationally accessible neighbors.

The three null models differ in whether the mutationally accessible variants are defined at the level of nucleotides, codons, or amino acids. In model 1, we define the set of possible mutant alleles as the set of all single-nucleotide mutations, and because each site has the same number of mutational neighbors (i.e., 3), the expected frequency of CpG mutants is simply the frequency of nucleotides found in CpG sites. In model 2, we assume that mutational neighbors that are synonymous or missense mutations are not likely to be chosen as the fittest neighbor, and so we exclude these. Finally, in model 3, we assume that – at any given site – only the amino acid state is relevant to fitness, so each possible amino acid state has the same chance of being the most fit, regardless of the number of nucleotide changes required to produce the state. Under this model, each amino acid change is assigned to be CpG-associated or not depending on whether or not the underlying nucleotide change in the ancestral codon would have occurred at a CpG site. Because our calculations (for all three models) are based on the reconstructed ancestral sequences, they naturally take into account the observed frequencies of CpG sites, amino acids, synonymous codons, and NNC GNN codon pairs in avian globin genes.

We note that for model 3 the structure of the genetic code renders all amino acid substitutions from an ancestral codon unambiguous with respect to whether they are CpG-associated, with exactly one exception. For the sequence TTC GNN, the first codon (which specifies Phe) can reach a neighbor (Leu) by either a CpG mutation (TTC GNN to TTR GNN) or a non-CpG mutation (TTC GNN to CTC GNN). We counted this as a CpG-associated change, which is conservative because it over-estimates the expected frequency of CpG changes, thereby making it more difficult to reject the null hypothesis.

### (f) Statistical tests for elevated frequency of CpG-associated changes

Having calculated the frequencies of CpG-associated changes under our null model, we then calculated the frequency of CpG-associated changes out of all observed adaptive mutations. Following [9], we note that because identical causative mutations sometimes contributed to adapation in several different high-altitude taxa it is useful to distinguish between the set of distinct mutations that contribute to adaptation, which we call “paths”, and the number of episodes of adaptation each such mutation contributed to, which we call “events”. We thus calculated the frequency of CpG-associated causative changes for both paths, where each distinct mutation is weighted equally, and for events, where each each mutation is weighted by the number of different adaptive episodes it contributed to. As described below, for the dataset in question we observe 10 different paths with 1 to 6 events per path. See Fig. 3 for all observed paths with two or more events.

**Figure 3.**
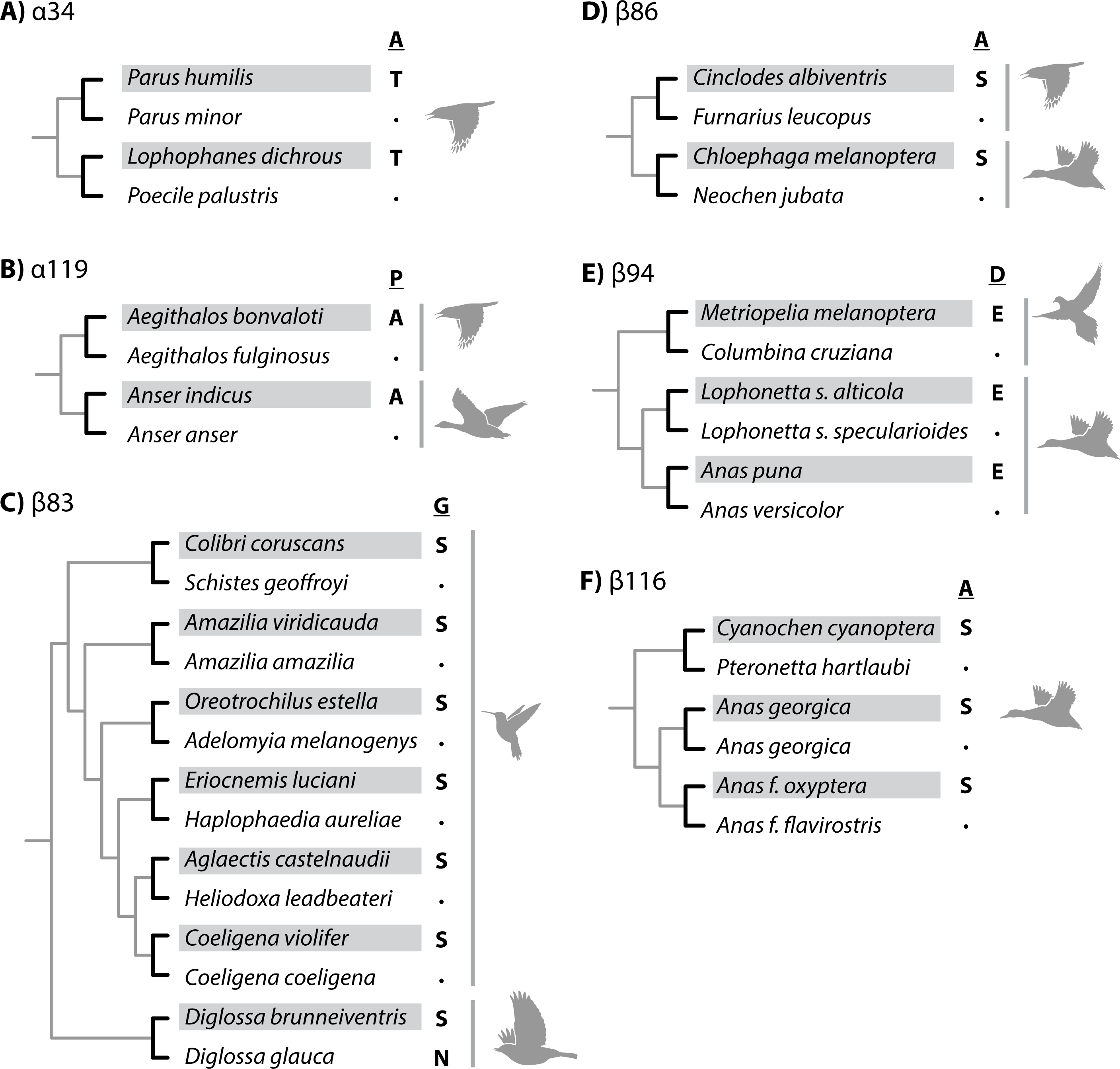
Phylogenetically replicated substitutions that contributed to derived increases in Hb-O_2_ affinity in different lineages of high-altitude birds. The select pairs of high- and low-altitude taxa are taken from the set of 70 taxa shown in Figure 1. In each case, the high-altitude taxon is denoted by shading and the inferred ancestral amino acid is shown as the reference state. A) Parallel substitutions at α34 in two high-altitude passerines from the Sino-Himalayan region. Paired comparisons between high- and low-altitude taxa involved ground tit, *Parus humilis* (high) vs. oriental tit, *P. minor* (low), and grey-crested tit, *Lophophanes dichrous* (high) vs. marsh tit, *Poecile palustris* (low)[26]. B) Parallel substitutions at α119 in high-altitude passerine and waterfowl species from the Sino-Himalayan region. Comparisons involved black-browed bushtit, *Aegithalos bonvaloti* (high) vs. sooty bushtit, *A. fulginosus* (low), and bar-headed goose, *Anser indicus* (high) vs. greylag goose, *Anser anser* (low)[26, 35, 36]. C) Parallel and convergent substitutions at β83 in high-altitude hummingbird and passerine species from the Andes. Comparisons involved sparkling violetear, *Colibri coruscans* (high) vs. wedge-billed hummingbird, *Schistes geoffroyi* (low), green-and-white hummingbird, *Amazilia viridicauda* (high) vs. amazilia hummingbird, *A. amazilia* (low), Andean hillstar, *Oreotrochilus estella* (high) vs. speckled hummingbird, *Adelomyia melanogenys* (low), sapphire-vented puffleg, *Eriocnemis luciani* (high) vs. greenish puffleg, *Haplophaedia aurelieae* (low), white-tufted sunbeam, *Aglaeactis castelnaudii* (high) vs. violet-fronted brilliant, *Heliodoxa leadbeateri* (low), violet-throated starfrontlet, *Coeligena violifer* (high) vs. bronzy inca, *C. coeligena* (low), and black-throated flowerpiercer, *Diglossa brunneiventris* (high) vs. deep-blue flowerpiercer, *Diglossa glauca* (low)[27, 29]. Note that the inferred ancestral state for β83 of hummingbirds is glycine (G), but the inferred ancestral state for this same site in *Diglossa* is asparagine (N). Thus, the βN83S substitution in *Diglossa brunneiventris* species represents a convergent substitution relative to the hummingbird clade. D) Parallel βA86S substitutions in high-altitude suboscine passerines and waterfowl species from the Andes. Comparisons involved cream-winged cinclodes, *Cinclodes albiventris* (high) vs. pale-legged hornero, *Furnarius leucopus* (low), and Andean goose, *Chloephaga melanoptera* (high) vs. Orinoco goose, *Neochen jubata* (low)[28, 29]. E) Parallel βD94E substitutions in high-altitude doves and waterfowl species from the Andes. Comparisons involved black-winged ground dove, *Metriopelia melanoptera* vs. croaking ground-dove, *Columbina cruziana*, Andean crested duck, *Lophonetta s. alticola* (high) vs. Patagonian crested duck, *L. s. specularioides* (low), and Puna teal, *Anas puna* (high) vs. silver teal, *A. versicolor* (low)[28, 29]. F) Replicated βA116S substitutions in high-altitude waterfowl. Comparisons involve Abyssinian blue-winged goose, *Cyanochen cyanoptera* (high) vs. Hartlaub’s duck, *Pteronetta hartlaubi* (low) and high-vs. low-altitude subspecies of yellow-billed pintail, *Anas georgica*, and sharp-winged teal, *Anas flavirostris*. In the latter two taxon pairs, the derived Ser-β116 variants do not have independent mutational origins in the two highland taxa; the allele-sharing is attributable to a history of introgressive hybridization [28].

After calculating the expected and observed frequencies of CpG-associated causative changes, we tested whether the observed frequency was significantly larger than the expected frequency our null model. However, these tests were somewhat different depending on whether we calculated the frequency at the level of paths or at the level of events. For paths, under our null hypothesis, each path is either CpG-associated or not CpG-associated independently of all other paths, and each path is CpG-associated with the same probability (as specified by one of the three null models). That is, under our null hypothesis, the number of CpG-associated paths is binomially distributed. We therefore evaluated this null hypothesis against the alternative hypothesis (that the frequency of CpG mutations among paths is greater than the null expectation) using a binomial test (binom.test in the R stats package).

Whereas paths are independent of one another, the causative mutations involved in different events may or may not have distinct mutational origins. Causative alleles that are shared between two or more high-altitude species may be identical-by-state (distinct mutational origins) or identical-by-descent (due to retention of ancestral polymorphism of introgressive hybridization). Thus, under the null hypothesis, different events belonging to the same path may not be independently CpG-associated. To account for this potential non-independence, we considered a null model in which the number of CpG-associated events is determined by randomly and independently assigning each observed path to be either CpG-associated or not with probabilities specified by one of the three null models. The frequency of CpG-associated events is then determined by weighting each path by the empirically observed number of events for that path. This choice of null is highly conservative because it increases the variability in the fraction of CpG-associated events relative to a more realistic but harder-to-specify null in which events with distinct mutational origins are treated as independent. To implement this randomization test for whether the number of CpG-associated events exceeds the null expectation we calculated the exact probability that the number of CpG events is greater than or equal to the observed number (custom R script, Dryad Digital Repository: https://datadryad.org/review?doi=doi:10.5061/dryad.2256f38).

### (g) Ancestral sequence reconstructions

We used alignments of avian globin sequences in conjunction with previously estimated phylogenetic relationships [40, 46] to reconstruct ancestral sequences using baseml, as implemented in PAML 4.7 [47]. We used the substitution model GTR+G in the ancestral sequence reconstructions of both α^*A*^- and β^*A*^-globin.

## 3. Results

### (a) Convergence in protein function

Given that theoretical and experimental results indicate that an increased Hb-O_2_ affinity can contribute to an enhancement of arterial O_2_ saturation and, hence, improved tissue O_2_ delivery under conditions of severe hypoxia [30-32], an obvious prediction is that derived increases in Hb-O_2_ affinity will have evolved repeatedly in avian taxa that have independently colonized extreme altitudes. We tested this prediction using phylogenetically independent comparisons involving 35 pairs of high- and low-altitude avian taxa (**Fig. 1**). The analysis revealed a striking elevational pattern of convergence, as the high-altitude taxon exhibited a higher Hb-O_2_ affinity in the overwhelming majority of pairwise comparisons. Phylogenetically independent comparisons revealed that highland natives generally have an increased Hb-O_2_ affinity relative to their lowland counterparts, a pattern consistent for both the major HbA isoform (Wilcoxon’s signed-rank test, *Z* = −4.6844, *P* < 0.0001, *N* = 35; **Fig. 2A**) and the minor HbD isoform (*Z* = −3.3144, *P* = 0.0009, *N* = 26; **Fig. 2B**). In all pairwise comparisons in which the high-altitude taxa exhibited significantly higher Hb-O_2_ affinities relative to the lowland taxa (*N* = 35 taxon pairs for HbA, *N* = 26 for HbD), the measured differences were entirely attributable to differences in intrinsic O_2_ affinity rather than differences in responsiveness to the inhibitory effects of Cl^-^ ions or IHP [26, 28, 29, 35]. The sample size is smaller for HbD because some species included in our dataset do not express this isoform [29, 39, 40].

### (b) Identification of causative mutations that contribute to convergent changes in Hb function

After documenting that high-altitude taxa have convergently evolved derived increases in Hb-O_2_ affinity, we used results of our previously published site-directed mutagenesis experiments to identify causative amino acid substitutions. Comparative sequence data for the set of 70 taxa revealed phylogenetically replicated replacements at numerous sites in the α^*A*^- and α^*D*^-globin genes (affecting HbA and HbD, respectively) and in the β^*A*^-globin gene (affecting both HbA and HbD) [26-29, 33, 35, 36]. Although we observed numerous parallel substitutions (i.e., independent changes from the same ancestral amino acid to the same derived amino acid) and convergent substitutions (i.e., independent changes from different ancestral amino acids to the same derived amino acid), functional data from native Hb variants and engineered, recombinant Hb mutants revealed that only a subset of replicated replacements actually contributed to convergent increases in Hb-O_2_ affinity in the different highland taxa. Overall, we identified a total of 22 affinity-enhancing replacements in 20 different high-altitude taxa (**Table 1**). This set of causative changes included two parallel replacements in α^*A*^-globin (αA34T and αP119A) and four parallel or convergent replacements in β^*A*^-globin (βG83S, βN83S, βA86S, βD94E, and βA116S)[26-29, 36](**Fig. 3**). Convergent increases in Hb-O_2_ affinity were largely attributable to nonreplicated replacements (divergent substitutions), suggesting that evolutionary increases in Hb-O_2_ affinity can be produced by amino acid replacements at numerous sites.

**Table 1.** Experimentally verified mutations (*, CpG mutations) that contributed to derived increases in Hb-O_2_ affinity in high-altitude birds.

| High-altitude taxon | Affinity-enhancing amino acid change | Nucleotide change | Reference |
| --- | --- | --- | --- |
| Bar-headed goose, *Anser indicus* | $\alpha 18$ G→S | G* to A | [36] |
| Grey-crested tit, *Lophophanes dichrous* | $\alpha 34$ A→T | G* to A | [26] |
| Ground tit, *Parus humilis* | $\alpha 34$ A→T | G* to A | [26] |
| Bar-headed goose, *Anser indicus* | $\alpha 63$ A→V | C* to T | [36] |
| Black-browed bushtit, *Aegithalos bonvaloti* | $\alpha 119$ P→A | C to G | [26] |
| Bar-headed goose, *Anser indicus* | $\alpha 119$ P→A | C to G | [36] |
| House wren, *Troglodytes aedon* | $\beta 55$ V→I | G* to A | [25] |
| Sparkling violetear, *Colubri coruscans* | $\beta 83$ G→S | G to A | [27, 29] |
| Green-and-white hummingbird, *Amazilia viridicauda* | $\beta 83$ G→S | G to A | [27, 29] |
| Andean hillstar, *Oreotrochilus estella* | $\beta 83$ G→S | G to A | [27, 29] |
| Sapphire-vented puffleg, *Eriocnemis luciani* | $\beta 83$ G→S | G to A | [27, 29] |
| White-tufted sunbeam, *Aglaeactis castelnaudii* | $\beta 83$ G→S | G to A | [27, 29] |
| Violet-thoated starfrontlet, *Coeligena violifer* | $\beta 83$ G→S | G to A | [27, 29] |
| Black-throated flowerpiercer, *Diglossa brunneiventris* | $\beta 83$ N→S | A to G | [29] |
| Cream-winged cinclodes, *Cinclodes albiventris* | $\beta 86$ A→S | G* to T | [29] |
| Andean goose, *Chloephaga melanoptera* | $\beta 86$ A→S | G* to T | [28] |
| Black-winged ground-dove, *Metriopelia melanoptera* | $\beta 94$ D→E | C to G | [29] |
| Andean crested duck, *Lophonetta specularioides alticola* | $\beta 94$ D→E | C to G | [28] |
| Puna teal, *Anas puna* | $\beta 94$ D→E | C to G | [28] |
| Yellow-billed pintail, *Anas georgica* | $\beta 116$ A→S | G* to T | [28] |
| Sharp-winged teal, *Anas flavirostris oxyptera* | $\beta 116$ A→S | G* to T | [28] |
| Abyssinian blue-winged goose, *Cyanochen cyanoptera* | $\beta 116$ A→S | G* to T | [28] |

### (c) Testing for a disproportionate contribution of CpG mutations to the adaptive convergence of Hb function

After experimentally identifying the causative substitutions that are responsible for (putatively adaptive) increases in Hb-O_2_ affinity in high-altitude birds, we then asked whether mutations at CpG sites made a disproportionate contribution to the observed changes in protein function. The use of site-directed mutagenesis experiments to test functional effects was restricted to the major HbA isoform (**Table 1**), so we focus attention on amino acid replacements in the α^*A*^- and β^*A*^- globin genes. Note, however, that the β^*A*^ *-*globin gene encodes β*-*type subunits of both HbA and HbD, so the effects of missense mutations in that gene are manifest in both isoforms.

The set of mutational changes in the α^*A*^ *-* and β^*A*^-globin genes can be described in several ways. There are 22 total substitution events that take place at 9 different sites via 10 different mutational paths. There is a difference between sites and paths because there are two different paths of change at site β83: there are parallel Gly→Ser replacements in six different high-altitude hummingbird species and there is a convergent Asn→Ser replacement in one species of high-altitude flowerpiercer (**Fig. 3**). The βG83S change, with 6 events, is the most highly repeated path. The distribution of events for all 10 paths is 6, 3, 3, 2, 2, 2, 1, 1, 1, and 1.

We calculated the expected frequency of CpG-associated changes under a null model in which the CpG status of a mutation is irrelevant to its probability of contributing to adaptation. Because this expectation varies slightly depending on assumptions, we calculated three separate null expectations (see Methods): (1) the frequency that a site in a globin gene is a CpG site, (2) the frequency that a missense mutation in a globin gene is a CpG mutation, and (3) the frequency of CpG-associated changes among the set of all one-mutant amino acid neighbors of a given ancestral sequence. The three models are very similar in giving a null expectation of 10 or 11 % CpG, as shown in Table 2. Thus, for the 10 independent mutational paths, we expect ∼1 CpG path, and for the 22 events, we expect ∼2 CpG events. As shown in Table 2, the observed number of CpG changes far exceeded the null expectations. Out of 22 events, 10 involve CpG mutations. Out of 10 paths, 6 involve a CpG mutation. The binomial test revealed that the observed frequency of CpG paths is significantly higher than the null expectation (**Table 2**). Likewise, the observed frequency of CpG mutations among events is significantly higher than the null expectation as determined by the randomization test that accounts for potential non-independence. These results remain qualitatively unchanged if we exclude βV55I from our set of causative mutations on the grounds that our initial discovery of this particular CpG path [25] motivated our investigation of CpG effects in the broader set of taxa. Leaving out βV55I and using model 1 (the most conservative), the probability of observing so many CpG-associated changes is *P*= 0.001 for paths (binomial test) and *P*= 0.033 for events (randomization test).

**Table 2.** Comparison of observed and expected numbers of CpG mutations that contribute to derived increases in Hb-O_2_ affinity in high-altitude species. See *Methods* for an explanation of the difference between mutational paths and events

| Model | f(expected) | CpG paths ($N = 10$) | | | CpG events ($N = 22$) | | |
| --- | --- | --- | --- | --- | --- | --- | --- |
| | | exp | obs | $P$ | exp | obs | $P$ |
| 1 | 0.109 | 1.1 | 6 | $2.4 \times 10^{-4}$ | 2.4 | 10 | 0.018 |
| 2 | 0.097 | 1.0 | 6 | $1.2 \times 10^{-4}$ | 2.1 | 10 | 0.013 |
| 3 | 0.096 | 1.0 | 6 | $1.2 \times 10^{-4}$ | 2.1 | 10 | 0.013 |

## 4. Discussion

### (a) A disproportionate role for CpG mutations in adaptation

Results of our comparative analysis of Hb function reveal that high-altitude birds have generally evolved increased Hb-O_2_ affinities relative to lowland sister taxa, consistent with earlier reports based on smaller subsets of the data presented here [26, 28, 29]. The striking pattern of convergence is consistent with the hypothesis that the elevational differences reflect a history of directional selection in high-altitude natives. This pattern of apparently adaptive convergence in protein function allowed us to address a key question: Are some types of amino acid mutation preferentially fixed? If so, to what extent is the observed substitution bias attributable to variation in rates of origin (mutation bias) – that is, variation among sites in rates of mutation to alleles that produce the beneficial change in phenotype? Our analysis of causative amino acid changes revealed that a disproportionate number of affinity-enhancing amino acid replacements were attributable to mutations at CpG dinucleotides, suggesting that mutation bias exerts an influence on patterns of adaptive substitution.

In each of the high-altitude bird species that evolved increased Hb-O_2_ affinities due to missense CpG mutations, there seems little reason to suppose that the causative amino acid mutations would have had larger selection coefficients (and, hence, higher fixation probabilities) than any number of other possible mutations that could have produced a similar increase in Hb-O_2_ affinity. However, if the rate of CpG mutation occurs at a higher rate than non-CpG mutations, then – in the absence of contributions from standing variation – the bias in mutation rate is expected to influence evolutionary outcomes in the same way as a commensurate bias in fixation probability [5, 7, 8].

### (b) Mutational independence of parallel substitutions

The majority of ‘repeated’ substitutions are authentic parallelisms, where the shared, derived amino acids in two or more high-altitude taxa have independent mutational origins [26, 28, 29]. However, in the case of the six repeated βG83S changes in high-altitude hummingbirds, we cannot rule out the possibility that some or most of the shared, derived Ser alleles in high-altitude species are identical-by-descent [27]. Even if the pattern of allele-sharing among species is partly attributable to incomplete lineage sorting or introgression, the pattern is still highly nonrandom with respect to altitude, suggesting a nonrandom sorting of ancestral polymorphism at β83 (or extensive introgressive hybridization). In other words, the observed pattern indicates that the derived Ser variant was repeatedly driven to fixation in different highland lineages [27]. Similarly, we previously documented that the shared, derived β116Ser variants in the high-altitude subspecies of *Anas georgica* and *Anas flavirostris* are identical-by-descent. In this case, the allele-sharing is attributable to introgressive hybridization, and results of population genomic analyses provide strong evidence for both highland taxa that the derived β116Ser variants increased in frequency under the influence of positive directional selection [28].

Thus, in the case of the repeated substitutions at β83 and β116, the derived amino acid variants do not necessarily have distinct mutational origins in different species, but the derived amino acid variants contributed to altitude-related increases in Hb-O_2_ affinity in each case, and this is what matters for the purposes of our tests. Using both binomial and randomization approaches, we tested the null hypothesis that the contributions of mutations to adaptive changes in Hb function is unrelated to whether they occur at CpG sites or non-CpG sites. Whether fixed alleles in different species are identical-by-descent is certainly relevant to questions about the prevalence of molecular parallelism and the causes of homoplasy [48, 49], but it has no bearing on our tests of whether CpG sites make disproportionate contributions to molecular adaptation. For example, in the case of the six βG83S substitutions in high-altitude hummingbirds, the ‘jackpot’ of six events in the same mutational path does not bias our conclusions because the possibility of such jackpot effects is incorporated into the design of the randomization test, which treats each observed mutational path as if it had a single mutational origin.

### (c) Conclusion: Mutation bias and substitution bias

We tested whether affinity-enhancing mutations in the Hbs of high-altitude birds are enriched for mutations at CpG dinucleotides relative to the frequency of CpG mutations among all possible missense mutations. The results summarized in Tables 1 and 2 indicate that a disproportionate number of causative amino acid replacements were attributable to CpG mutations, suggesting that mutation bias can influence outcomes of molecular adaptation. Moreover, if methylated CpG sites have higher-than-average rates of point mutation, we hypothesize that any given set of adaptive substitutions should be enriched for changes at such sites.

## Data availability

The supplementary data consists of (1) nucleotide and protein sequence alignments for the α^*A*^-, α^*D*^-, and β^*A*^-globin genes and (2) scripts used to perform calculations. All data are available in the Dryad Digital Repository: https://datadryad.org/review?doi=doi:10.5061/dryad.2256f38.

## Author contributions

JFS, DMM, and AS conceived the study; JFS, DMM, and AS drafted the manuscript; CN and AVS collected experimental data; JFS, AVS, and CCW analyzed experimental data; DMM and AS designed and implemented the statistical analysis of CpG effects; AS assembled and curated the supplementary data package.

## Acknowledgement

The identification of any specific commercial products is for the purpose of specifying a protocol, and does not imply a recommendation or endorsement by the National Institute of Standards and Technology.

